# Suppression of a dsRNA-induced plant immunity pathway by viral movement protein

**DOI:** 10.1101/2021.10.30.466425

**Authors:** Caiping Huang, Ana Rocio Sede, Jerome Mutterer, Emmanuel Boutant, Manfred Heinlein

**Author notes:** Current address: UMR 7021, CNRS, Laboratoire de Bioimagerie et Pathologies, Université de Strasbourg, Faculté de Pharmacie, 67400 Illkirch, France.

## Abstract

The virome of plants is dominated by RNA viruses ^1^ and several of these cause devastating diseases in cultivated plants leading to global crop losses ^2^. To infect plants, RNA viruses engage in complex interactions with compatible plant hosts. In cells at the spreading infection front, RNA viruses replicate their genome through double-stranded RNA (dsRNA) intermediates and interact with cellular transport processes to achieve cell-to-cell movement of replicated genome copies through cell wall channels called plasmodesmata (PD) ^3^. In order to propagate, viruses also must overcome host defense responses. In addition to triggering the antiviral RNA silencing response, RNA virus infection also elicits pattern-triggered immunity (PTI) ^4^ whereby dsRNA, a hallmark of virus replication, acts as an important elicitor ^5^. This innate antiviral immune response is also triggered when dsRNA is applied externally and does not require sequence homology to the virus ^5^. However, the mechanism by which PTI restricts virus infection is not known. Here, we show that dsRNA inhibits the progression of virus infection by triggering callose deposition at plasmodesmata and the inhibition of transport through these cell-to-cell communication channels. The dsRNA-induced signaling pathway leading to callose deposition is independent of ROS production and thus distinguished from pathways triggered by bacterial and fungal elicitors. The dsRNA-induced host response at plasmodesmata is suppressed by the *Tobacco mosaic virus* movement protein (MP). Thus, the virus uses MP to inhibit innate dsRNA-induced immunity at plasmodesmata, which could be a general strategy of phytoviruses to overcome plant defenses and spread infection.

## Text

RNA silencing is the most important antiviral host response in plants. It involves host DICER-LIKE enzymes that cleave viral dsRNA into small interfering RNAs (siRNAs). These viral siRNAs associate with ARGONAUTE proteins in RNA-induced silencing complexes (RISCs) to guide the sequence-specific degradation and translational suppression of the viral RNA genome. To control this antiviral response, viruses have evolved specific effector proteins that interfere with the RNA silencing pathway at distinct steps ^6^. Unlike RNA silencing, dsRNA-induced PTI is independent of dsRNA sequence and also activated by non-viral dsRNA, for example by the synthetic dsRNA analog polyinosinic-polycytidilic acid [poly (I:C)]. Similar to virus replication, treatment of *Arabidopsis thaliana* plants with poly (I:C) elicits strong antiviral defense along with activating typical PTI markers, such as mitogen-activated protein kinase (MPK), ethylene production, seedling root growth inhibition, and marker gene expression ^4,5^.

To discover how dsRNA-induced PTI inhibits RNA virus infection, we visualized the effect of poly(I:C) treatment on local infections of *Nicotiana benthamiana* plants using *Tobacco mosaic virus* tagged with green fluorescent protein (TMV-GFP). The TMV-GFP infection sites were smaller at 7 days post inoculation (dpi) in plants treated with poly(I:C) or with a bacterial PTI elicitor derived from flagellin (flg22) than in control plants treated with buffer (**Fig. 1**). The treatments did not cause a significant change in GFP fluorescence (**Fig. 1**) and had no bulk effect on viral RNA accumulation in the infected cells (**Fig. S1**). The smaller infection sites in treated leaves suggested that the immunity is linked to reduced cell-to-cell movement of the virus.

**Fig. 1.**
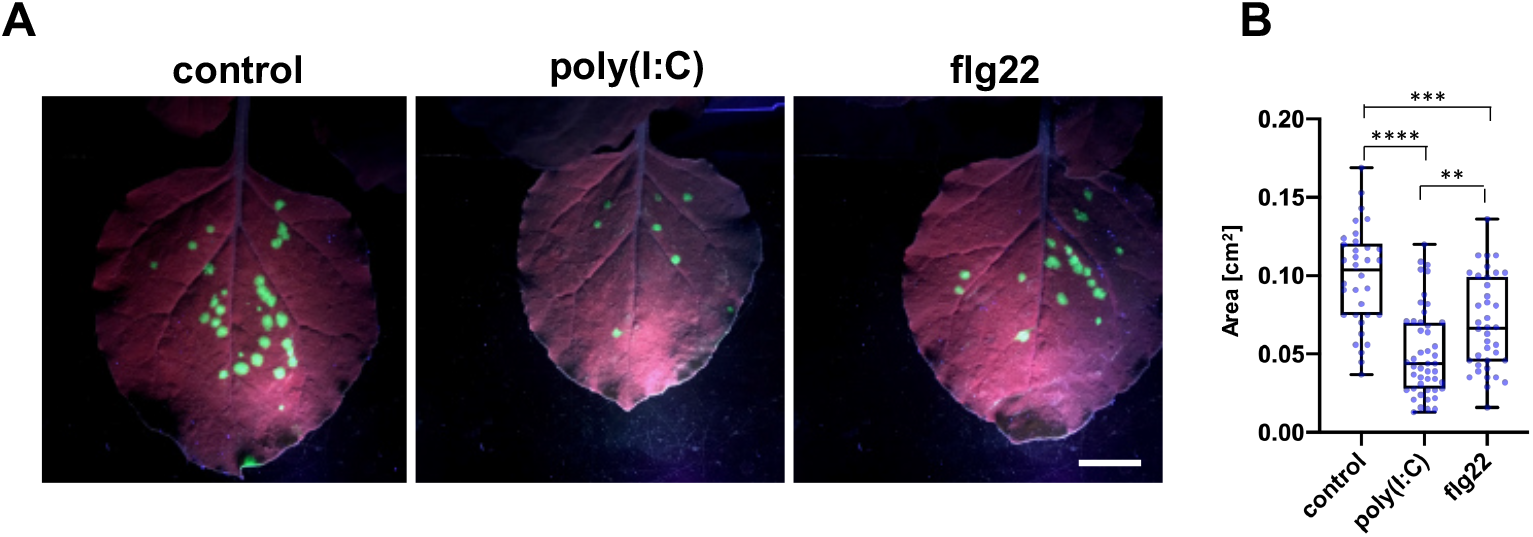
dsRNA causes inhibition of virus movement in *N. benthamiana*. (**A**) TMV-GFP infection sites in *N. benthamiana* leaves at 7 dpi with the virus together with either water (control), 0.5 µg/µl (≈1 µM) poly(I:C), or 1 µM flg22. (**B**) Sizes of individual infection sites measured in 10 leaf samples per treatment. Scale bar, 2 cm. Two-tailed Mann-Whitney test; ****, p <0,0001; ***, p <0,001; **, p <0,01.

Virus movement occurs through plasmodesmata (PD). We thus hypothesized that dsRNA inhibits virus movement by causing PD closure. Treatment of *N. benthamiana* plants with poly(I:C), flg22, or water, and quantification of PD-associated callose by *in vivo* aniline blue staining revealed that both poly(I:C) and flg22 trigger an increase in PD-associated callose levels in a concentration-dependent manner (**Fig. 2A** and **2B**). In agreement with this observation, treated tissues showed reduced PD permeability as determined by a GFP mobility assay (**Fig. 2C** and **2D**). As previously noted in Arabidopsis ^5^, poly(I:C) triggered MPK activation but the level of activation is significantly weaker than the activation observed with flg22 (**Fig. 2E**). Treated leaves exhibited the induction of *N. benthamiana* defense marker genes, such as genes encoding BOTRYTIS INDUCED KINASE1 (BIK1), PATHOGENESIS-RELATED PROTEIN 2 (PR2), NADPH/RESPIRATORY BURST OXIDASE PROTEIN B (RbohB) and ENHANCED DISEASE SUSCEPTIBILITY 1 (EDS1), but not the gene for BRASSINOSTEROID INSENSITIVE 1 (BRI1) (**Fig. S2**).

**Fig. 2.**
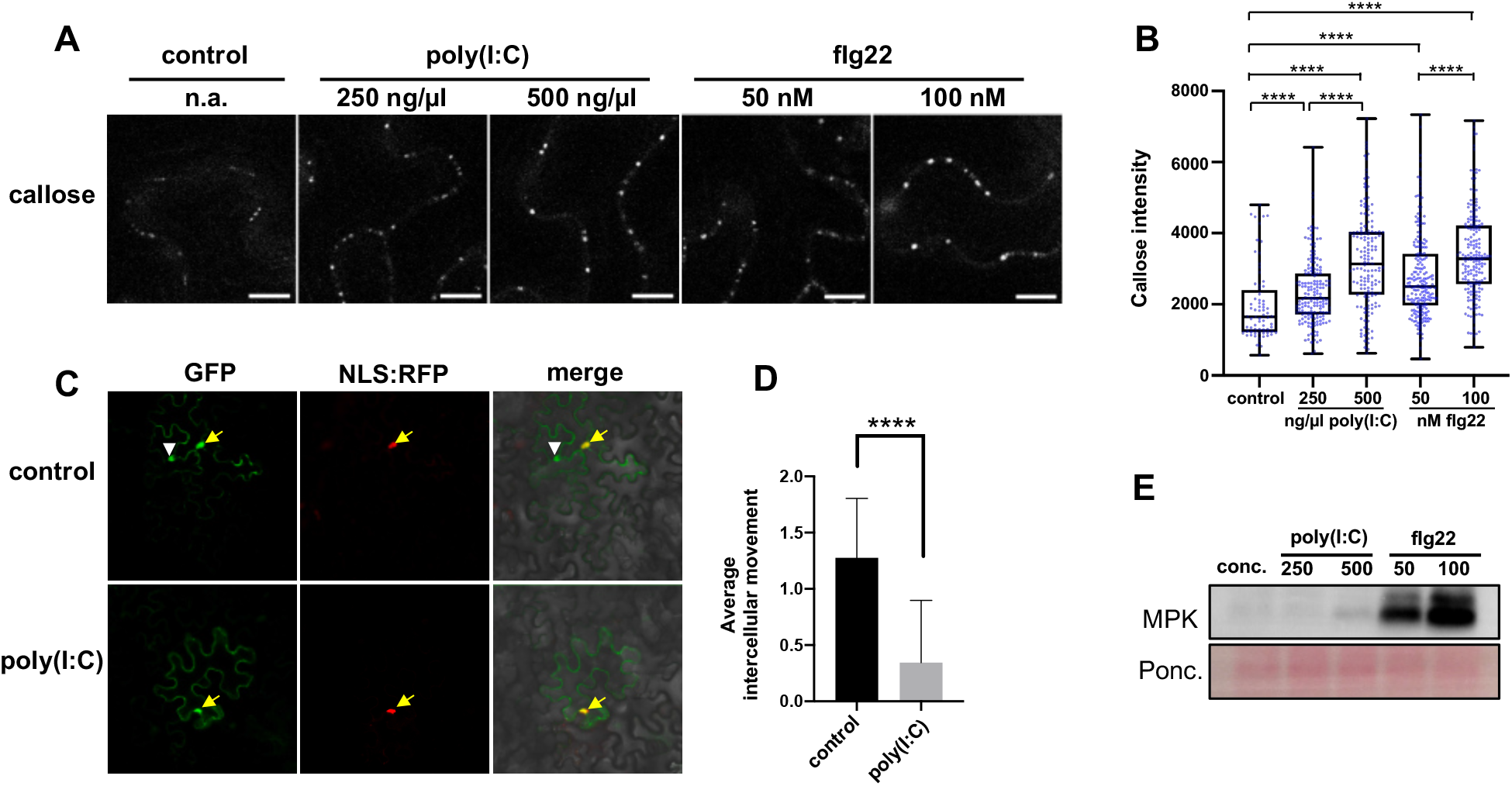
dsRNA induces PD closure. (**A**) and (**B**) Treatment of *N. benthamiana* leaves with poly(I:C) or flg22 induces increased callose deposition at PD within 30 minutes. (**A**) Callose fluorescence at PD seen upon aniline blue staining. Scale bar, 10 µm. (**B**) Relative callose content in individual PD (blue dots). Two-tailed Mann-Whitney test; ****, p = <0,0001. (**C**) and (**D**) GFP mobility assay. (**C)** GFP shows a nucleocytoplasmic distribution (yellow arrow) and its movement from the expressing cell (marked by co-expression with cell-autonomous NLS:RFP) is evident by appearance of green fluorescence in the nuclei of adjacent cells (white arrowhead). Scale bar, 50 µm. (**D)** Quantification of GFP movement (n = 30). Two-tailed Mann-Whitney test; ****, p = <0,0001. (**E**) Low level of MPK activation by poly(I:C) relative to flg22.

To determine how dsRNA elicits the deposition of callose at PD, we turned our attention to Arabidopsis. Treatment of *A. thaliana* Col-0 plants with poly(I:C) led to increases in PD-associated callose levels similar to those in *N. benthamiana* (**Fig. 3A**). These treatments confirmed the ability of poly(I:C) to activate MPK and to inhibit seedling growth (**Fig. S3, A-C**), as previously reported ^5^. Interestingly, dsRNA-induced callose deposition was strongly inhibited in *bik1 pbl1* plants (**Fig. 3B**), which are deficient in the receptor-like cytoplasmic kinase (RLCK) BIK1 and its homolog PBS1-LIKE KINASE1 (PBL1). BIK1 is a central component of PTI signaling that integrates signals from multiple pathogen-recognition receptors (PRRs), as shown by its direct interaction with FLS2, EFR, PEPRs, and CERK1 ^7–9^ and its ability to phosphorylate and activate downstream targets, such as the NADPH Oxidase RBOHD ^10^. BIK1 and PBL1 have additive effects; the *bik1 pbl1* double mutant was shown to strongly inhibit PAMP-induced defense responses ^8^. In addition to the defect in dsRNA-induced callose deposition, *bik1 pbl1* plants are also deficient in poly(I:C)-induced MPK activation and growth inhibition (**Fig. S3, D-F**). These findings establish a role of BIK1/PBL1 in the innate immunity triggered by dsRNA that leads to callose deposition at PD.

**Fig. 3.**
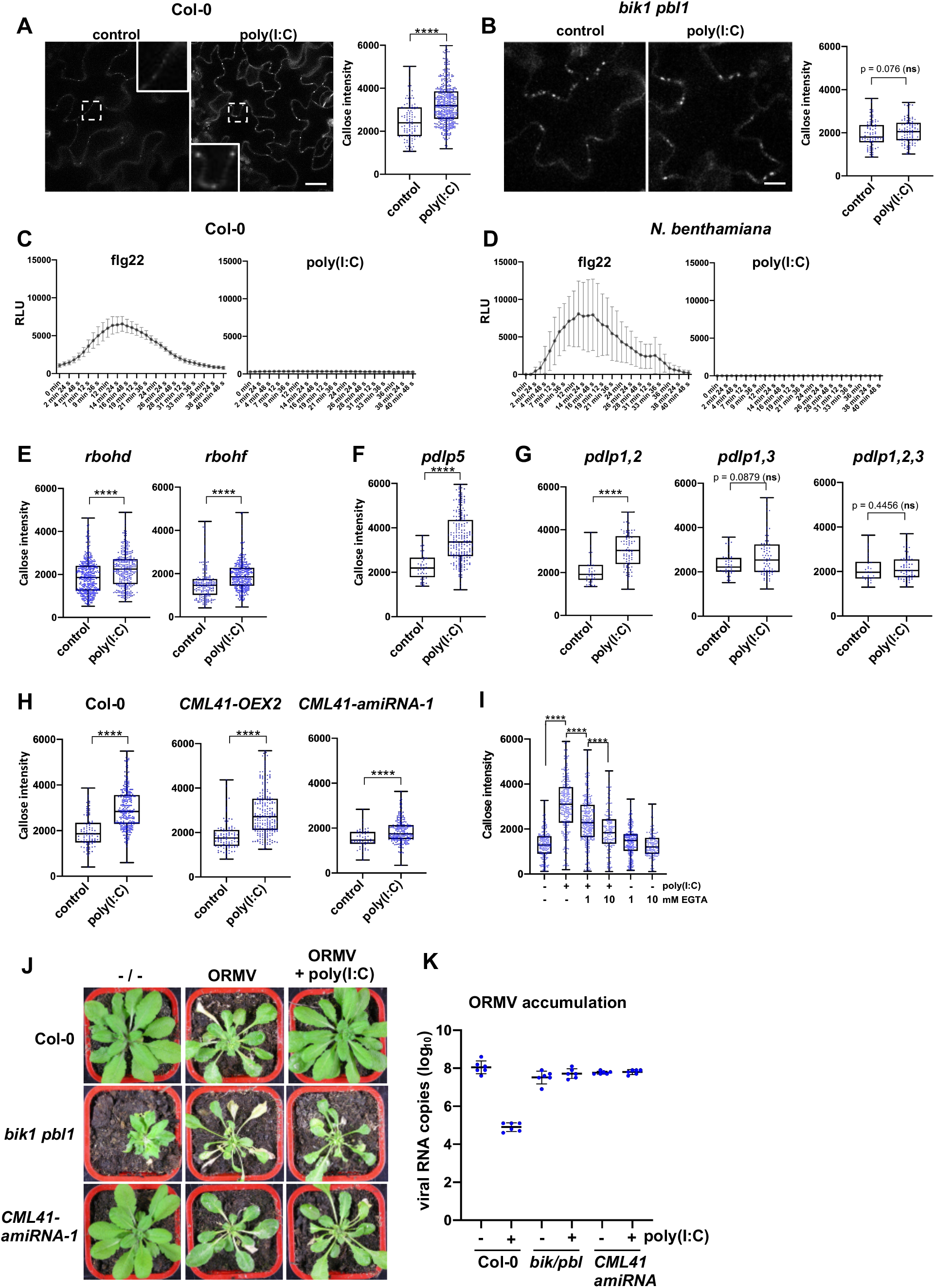
dsRNA-induced signaling in Arabidopsis. (**A**) poly(I:C) treatment causes callose deposition at PD. Inlays show enlargements of the areas within the dashed boxes. Scale bar, 20 µm. (**B**) dsRNA-induced callose deposition in PD is inhibited in the *bik1 pbl1* mutant. Scale bar, 10 µm. (**C**) and (**D**) unlike flg22, poly(I:C) does not induced any ROS production in Arabidopsis (**C**) and *N. benthamiana* (**D**). (**E**) dsRNA-induced callose deposition is not affected in *rbohd* or *rbohf* mutants. (**F**) and (**G**) Analysis of *pdlp* mutants. dsRNA-induced callose deposition is independent of PDLP5 but is inhibited in the *pdlp1,2,3* triple mutant. (**H**) Inhibition of dsRNA-induced callose deposition in *CML41-amiRNA-1*-expressing plants. (**I**) dsRNA-induced callose deposition is reduced in the presence of EGTA. (**J**) and (**K**) Symptoms (J) and viral RNA accumulation (K) in wild type plants and mutants that were inoculated with ORMV in the presence and absence of poly(I:C). Unlike in wild type plants (Col-0) the antiviral effect of dsRNA is lost in *bik1 pbl1* mutants and *CML41-amiRNA-1* expressing plants. Two-tailed Mann-Whitney test; ****, p <0,0001.

To further determine the signaling pathway downstream of BIK1, we examined whether the treatment of plants with poly(I:C) leads to the production of reactive oxygen species (ROS). To our surprise, neither the treatment of Arabidopsis Col-0 plants (**Fig. 3C**) nor the treatment of *N. benthamiana* plants (**Fig. 3D**) with poly(I:C) led to the production of ROS. By contrast, strong responses were recorded in both plant species upon treatment with the flg22 elicitor. These observations indicate that dsRNA-induced callose deposition and the dsRNA-based perception of viruses are independent of ROS. Confirming this result, *rbohd* and *rbohf* mutants deficient for the major ROS producing NADPH oxidases RBOHD and RBOHF ^11^ responded like wild-type Col-0 plants to the presence of dsRNA, showing induced callose deposition at PD (**Fig. 3E**).

To determine the signaling pathway induced by dsRNA, additional mutants were tested. We started with Arabidopsis mutants deficient in the PD-localized proteins (PDLPs), which are a family of eight proteins that dynamically regulate PD. For example, PDLP5 mediates salicylic acid (SA)-induced callose deposition and PD closure, which is required for plant resistance to bacterial pathogens ^12,13^. *pdlp5* mutant plants showed strong callose deposition at PD upon poly(I:C) treatment (**Fig. 3F**), indicating that dsRNA-induced callose deposition is independent of PDLP5 and, thus, of a potential SA response mediated through PDLP5. *pdlp1 pdlp2* double mutant plants also showed normal callose deposition. In contrast, *pdlp1 pdlp3* double mutant and *pdlp1 pdlp2 pdlp3* triple mutant plants were unable to significantly increase PD-associated callose levels in response to poly(I:C) (**Fig. 3G**). The involvement of PDLP1 and/or PDLP3 is consistent with their redundant roles in callose deposition at PD ^14^ and also in callose deposition within haustoria formed in response to infection by mildew fungus ^15^.

We found that dsRNA-induced callose deposition at PD also depends on the Ca^2+^-binding, PD-localized CALMODULIN-LIKE protein 41 (CML41). This protein was shown to mediate rapid callose deposition at PD associated with decreased PD permeability following flg22 treatment ^16^. Plants of a *CML41* overexpressing transgenic line (*CML41-OEX-2*) showed increases in PD-associated callose upon poly(I:C) treatment, like non-transgenic Col-0 controls. By contrast, plants in which *CML41* is downregulated by an artificial miRNA (*CML41-amiRNA-1*) showed a decreased response (**Fig. 3H**). Consistent with the role of CML41 in the callose deposition response to dsRNA, the permeability of PD was previously shown to be sensitive to cytosolic Ca^2+^ concentrations ^17,18^. To test the role of Ca^2+^ in dsRNA-triggered innate immunity, we treated plants with poly(I:C) together with EGTA, a Ca^2+^- chelating molecule. EGTA reduced the level of callose induced at PD after dsRNA treatment (**Fig. 3I**). This effect was EGTA concentration-dependent and indicates a role of Ca^2+^ in dsRNA-triggered PD regulation. Together, these results suggest a role for CML41 and Ca^2+^ in the dsRNA-induced defense response at PD.

We previously showed that poly(I:C) treatment protects Arabidopsis plants against infection by *Oilseed rape mosaic virus* (ORMV) ^5^. If BIK1 and CML41 mediate dsRNA-induced PD closure, we hypothesized that this antiviral defense should be weakened in *bik1 pbl1* mutant plants and in the *CML41 amiRNA-1* plants. Whereas poly(I:C) treatment prevented symptoms at 28 dpi and resulted in a strongly reduced virus titer in wild-type Col-0 plants (as previously reported) ^5^, both the *bik1 pbl1* and *CML41 amiRNA-1* plants showed symptoms and accumulated high virus levels in poly(I:C)-treated plants, like in ORMV-infected but untreated controls (**Fig. 3J** and **3K**). Ablation of virus-inoculated leaves from plants at different times after inoculation showed that the time required for the virus to exit the inoculated leaf and to cause systemic infection was 3 days in Col-0 plants. By contrast, this time was reduced to 24 hours in *bik pbl1* mutants and *CMLl41-amiRNA-1* plants (**Fig S4, A and B**). Taken together, these findings show that dsRNA-induced callose deposition at PD correlates with antiviral plant defense at the level of virus movement. Furthermore, the experiments reveal a signaling pathway that requires BIK1/PBL1, CML41 and Ca^2+^, as well as PDLP1 and PDLP3 for callose deposition. This dsRNA-induced callose deposition is independent of ROS and thereby distinguished from innate immunity pathways that close PD in the presence of fungal and bacterial elicitors ^10,19^.

The plant-pathogen arms race causes pathogens to evolve effectors that overcome host defenses. Therefore, we wondered whether the movement protein (MP) of TMV facilitates virus movement by suppressing the poly(I:C)-induced plant defense response at PD. To address this question, we divided the local infection site into different zones (**Fig. 4A**): zone I ahead of infection and without MP, zone II at the virus front where MP facilitates virus movement, zone III behind the infection front, and zone IV, which is the center of the infection site where MP is no longer expressed. Zones II-IV continuously accumulate dsRNA in distinct replication complexes that also produce MP (**Fig. S5**). Aniline blue staining demonstrates high PD-associated callose levels within and around the infection site (**Fig. 4B**). However, cells in zone II and zone III, where virus cell-to-cell movement is associated with a transient activity of MP in increasing the PD size exclusion limit ^20^, exhibit a marked reduction in PD-associated callose levels as compared to cells in zone I (ahead of infection) and zone IV (center of infection) (**Fig. 4B** and **C**). The low level of PD-associated callose in cells at the virus front (zone II) is consistent with the ability of MP to transiently suppress dsRNA-induced immunity at PD.

**Fig. 4.**
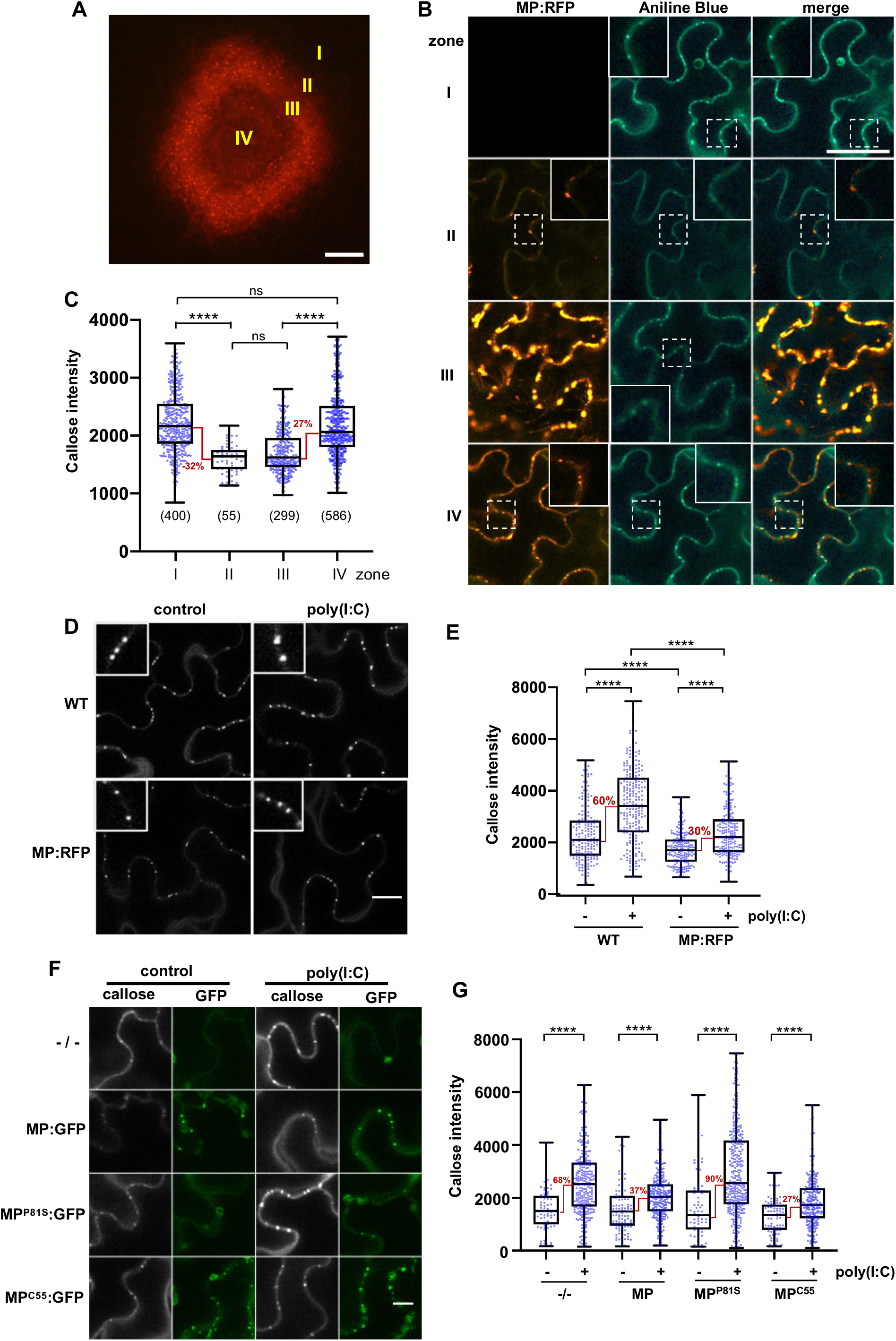
Suppression of dsRNA-induced immunity by MP. (**A**) Local site of infection by TMV-MP:RFP in *N. benthamiana*. Different zones ahead of infection (zone 1), at the infection front (zone II), behind, the infection front (zone III) and in the center of infection (zone IV) are indicated. Scale bar, 200 µm. (**B**) Pattern of MP:RFP and callose (Aniline Blue) accumulation in the different zones. Inlays show magnifications of the image areas highlighted by dashed boxes. Scale bar, 40 µm. (**C**) Quantification of PD callose in the different zones. Two-tailed Mann-Whitney test; ****, p <0,0001; ns, p>0.05 (**D**) and (**E**) Inhibition of dsRNA-induced callose deposition in MP:RFP-transgenic plants. Scale bar, 20 µm; Two-tailed Mann-Whitney test; ****, p <0,0001. (**F**) and (**G**) Inhibition of dsRNA-induced PD callose deposition by transiently expressed MP:GFP. The inhibitory effect is abolished by a single amino acid exchange mutation in MP (P81S) that inhibits its ability to target PD and to function in virus movement. Functional MP with C-terminal deletion of 55 amino acids (C55) targets PD and also inhibits dsRNA-induced callose deposition like wildtype MP. Scale bar, 10 µm. Two-tailed Mann-Whitney test; ****, p <0,0001.

To test this hypothesis, we examined whether MP suppresses the poly(I:C)-induced callose deposition at PD in the absence of viral infection. Transgenic *N. benthamiana* plants that stably express MP:RFP have the ability to complement a MP-deficient TMV mutant for movement, thus indicating that the MP:RFP in these plants is at least partially active (**Fig. S6**). Treatment of such plants with poly(I:C) showed a 50% lower induction of callose deposition at PD as compared to wild type plants (**Fig. 4D** and **E**). The induction of callose by poly(I:C) was also reduced upon transient expression of MP:GFP (**Fig. 4F** and **G**). Importantly, the same effect was observed with MP^C55^:GFP. This mutant MP lacks 55 amino acids from the C-terminus but still accumulates at PD and is functional in TMV movement ^21^. By contrast, dysfunctional MP^P81S^ carrying a P to S substitution at amino acid position 81, fails to target PD ^22^ and does not interfere with poly(I:C)-induced callose deposition. These experiments show that the TMV MP has the capacity to significantly reduce the dsRNA-induced callose deposition at PD, and that this inhibition of callose deposition requires a MP that can facilitate virus movement.

Replication of an MP-deficient TMV replicon was previously shown to induce callose deposition at PD ^23^, but the viral molecules inducing this deposition remained obscure. Consistent with findings that viruses induce innate immunity ^4^ and that dsRNA is a potent PAMP in plants ^5^, we show here that dsRNA-induced immunity leads to PD callose deposition and closure. The required signaling pathway involves BIK1/PBL1, CML41, Ca^2+^, and PDLP1,2,3 but not ROS production or PDLP5. Thus, although RBOHD and ROS production are a hallmark of PTI ^24^ and both ROS and PDLP5-mediated SA signaling play a role in PD regulation ^25^, other pathways also exist. The lack of ROS production distinguishes virus/dsRNA-induced signaling from the ROS-associated responses induced by other pathogens. The PD-localized CML41 protein was previously shown to participate in flg22-triggered, but not chitin-triggered PD callose deposition ^16^. Thus, although differing in upstream components, dsRNA and flg22-induced signaling act on shared PD-associated regulatory components. The absence of ROS signaling in dsRNA-induced PD regulation could reflect this specific elicitor type or its location of perception. Bacterial and fungal PAMPs are released in the apoplast and perceived by PRRs at the plasma membrane (PM), while viral dsRNA formation and perception may occur at intracellular membranes where viruses replicate. RIG-I-like receptors (RLRs) including RIG-I, MDA5 and LGP2 detect the presence of dsRNA in animals ^26^ and provide examples of dsRNA perception in the cytoplasm. Plant viruses like TMV and ORMV replicate in association with punctate sites at the cortical endoplasmic reticulum (ER) that may represent ER:PM contact sites that also occur at PD and may allow interaction of the viral replication complexes with PM-localized signaling proteins. Importantly, dsRNA-induced innate immunity is unaffected by mutations in dsRNA binding DICER-LIKE (DCL) proteins, which excludes these proteins as the dsRNA receptors for PTI and shows that dsRNA silencing and dsRNA-induced innate immunity require different protein machineries ^5^.

TMV and its MP represent the paradigm model for virus movement ^27^. The ability of MP to reduce poly(I:C)-induced callose deposition at PD is consistent with previous findings suggesting that viral MPs operate callose-degrading enzymes or PD structural components to increase the PD size exclusion limit ^28–30^. Our results widen this model by suggesting that viral MPs may act through interaction with components of the dsRNA-induced immunity pathway leading to PD closure. Identifying the PTI dsRNA receptor and also the components through which MP reduces the dsRNA-induced PTI response at PD will be the challenge for future studies.

## Acknowledgements

We thank Cyril Zipfel, Christine Faulkner, Matthew Gilliham, and Sacco de Vries for providing seeds of Arabidopsis mutants and Todd Blevins for text editing.

## Methods

### Plant materials and growth conditions

*N.benthamiana* and *A.thaliana* plants were grown from seeds in soil with 16h/8h light/dark periods at +22 °C/+18 °C. MP:RFP-transgenic *N.benthamiana* plants were produced by leaf disk transformation ^31^ using plasmid pH7-MP:RFP ^32^. *N. benthamiana* plants expressing GFP fused *Flock house virus* B2 protein have been described previously ^33^.

### Virus inoculation

cDNA constructs for TMV-MP:RFP ^34^, TMV-GFP ^35^, and TMVΔMP-GFP ^36^ have been described previously. *N. benthamiana* plants were mechanically inoculated the presence of an abrasive (Celite®545) with infectious RNA *in vitro*-transcribed from these constructs. Arabidopsis plants were inoculated with purified ORMV virions ^37^.

### Analysis of virus infection in the presence of elicitors

To test the effect of elicitors on TMV-GFP infection in *N. benthamiana*, the inoculum with infectious viral RNA was mixed with specific elicitor to contain a final concentration of 0.5 µg/μl (equals ∼1 µM) poly(I:C) or 1uM flg22. Infection sites on the inoculated leaves were imaged at 7 days post inoculation (dpi) using a hand-held camera and UV lamp (BLAK RAY B-100AP; UVP Inc., Upland, California) in the presence of a ruler for size normalization. The areas of infection sites in each leaf were measured with Image J software upon selection of infection site as regions of interest using fluorescence thresholding and the wand tracing tool, and by setting the scale according to the ruler.

To test the effect of elicitors on ORMV infection in Arabidopsis, 4 µl of elicitor solution (10 mg/ ml poly(I:C) or 10 µM flg22) or 4 µl PBS were placed on rosette leaves of 3 weeks old Arabidopsis wildtype or mutant Col-0 plants. A volume of 2.5 µl of a 20 ng /µl solution of purified ORMV virions was placed on the same leaves. Subsequently, the leaves were gently rubbed in the presence of celite as abrasive. Immediately after treatment, remaining elicitors, buffers and virions were washed off the leaf surface. Symptoms were analysed at 28 dpi. At the same time young, systemic leaves were sampled for analysis of virus accumulation by quantitative Taqman RT-PCR using previously described methods ^5,38^.

### Analysis of differential gene expression measurement by RT-qPCR

*N. Benthamiana* leaf discs were excised with a cork borer and incubated overnight in 12-well plates containing 600 μl deionized, ultra-pure water. The leaf disks were washed several times with water and then incubated with elicitor (1 μM flg22, 0.5 μg/μl poly(I:C), or water as control) for 3 hours. After washing the discs with deionized, ultra-pure water three times, samples were ground to a fine powder in liquid nitrogen and total RNA was extracted by TRIzol™ reagent according to the protocol of the manufacturer. 2 ug of RNA were reverse transcribed using a reverse transcription kit (GoScript™ Reverse Transcription System, Promega). The abundance of specific transcript was measured by probing 1 μL cDNA by quantitative real-time PCR in a total volume of 10 µl containg 5 μL SYBR-green master mix (Roche), 0.5 μM forward and reverse primer and water. PCR was performed in a Lightcycler480 (Roche) using a temperature regime consisting of 5 minutes at 95°C followed by 45 cycles at 95°C for 10 seconds, 60°C for 15 seconds, and 72°C for 15 seconds, and ending with a cycle of 95°C for 5 seconds, 55°C for 60 seconds, 95°C for continuous time until final cooling to 40°C for 30 seconds. The threshold cycle (CT) values were normalized to CT-values obtained for housekeeping genes, and used to calculate the relative expression levels, the mean values and standard errors (SE). Each mean value represents the analysis of three independent replicate samples, each measured by three technical replicates. Primers are listed in Table S1.

### Agroinfiltration

Plasmids for expression of MP:GFP, MP^C55^:GFP, and MP^P81S^:GFP were created by Gateway cloning and have been described previously ^32,39,40^.

For transient expression of the fluorescent fusion proteins, cultures of *A. tumefaciens* bacteria carrying these plasmids were harvested by centrifugation, resuspended in infiltration medium (10 mM MES, 10 mM MgCl_2_, 200 μM acetosyringone; pH 5.5) to a final optical density at 600 nm (OD_600_) of 0.1 (unless stated differently), and infiltrated into the abaxial side of the leaf using a syringe without a needle. Leaves were observed by confocal microscopy at 48 hours after agroinfiltration. For GFP mobility assays, we used Agrobacteria that were co-transformed with binary vectors for expression of GFP together with the cell-autonomous nuclear protein NLS:RFP (pB7-NLS:M2CP:RFP; this vector was created by recombining pZeo-NLS:MS2CP ^40^ with expression vector pH7RWG2). Before infiltration, the diluted culture (OD_600_ = 0.1) was further diluted 1: 1000 or 1:10000 to ensure that both proteins will be expressed in only few cells of the leaf. GFP movement was evaluated by counting the radial cell layers into which GFP has moved away from the infiltrated cell (marked by red fluorescent nucleus).

### Imaging

For all other imaging a Zeiss LSM 780 confocal laser scanning microscope with ZEN 2.3 software (Carl Zeiss, Jean, Germany) was used. Excitation / emission wavelengths were 405 nm / 475-525 nm for aniline blue, 488 nm / 500-525 nm for GFP, and 561 nm / 560-610nm for RFP.

### Callose staining

Leaf disks were excised with a cork borer and placed into wells of 12-well culture plates containing 1 ml water and incubated overnight under conditions at which the plants were raised. The leaf discs were washed several times with water before use. For callose staining, individual leaf disks were placed on microscope slides and covered with a coverslip fixed with tape. A mixture of 4 µl poly(I:C) (25 µg/ul in water) or 4 µl water as control added to 196 µl of 1% aniline blue solution (in 50 mM potassium phosphate buffer (pH 8.0)) was soaked into the space between the glass slide and coverslip. The glass slide with the sample was evacuated for 1-2 minutes (< 0.8 pa) in a vacuum desiccator followed by slow release of the pressure. Aniline blue fluorescence was imaged 30 minutes after dsRNA or control treatment using a Zeiss LSM 780 confocal laser scanning microscope with ZEN 2.3 software (Carl Zeiss, Jean, Germany) and using a 405 nm diode laser for excitation and filtering the emission at 475-525 nm. 8-bit Images acquired with a 40× 1.3 N.A. Plan Neofluar objective with oil immersion were analyzed with ImageJ software (http://rsbweb.nih.gov/ij/) using the plug-in *calloseQuant*, which after setting few parameters localizes fluorescent callose spots and quantifies callose fluorescence intensity of each spot automatically ^41^. This plugin is available at https://raw.githubusercontent.com/mutterer/callose/main/calloseQuant_.ijm. Callose spots were measured in 1-3 images taken from each leaf disk. For each genotype or condition, three leaf discs from three different plants were evaluated. To control for normal poly(I:C) treatment and callose staining conditions, samples of Arabidopsis mutant were always analyzed in parallel to samples from the Col-0 wild-type. The distribution of pooled fluorescence intensities obtained for the specific genotype or treatment condition is shown in boxplots. Individual fluorescence intensities that occurred as outlyers from the general distribution of fluorescence intensities (<1%) in the sample were excluded from boxplots and statistical analysis.

### Analysis of MPK activation

Leaf disks from 4-week-old *A.thaliana* or *N. Benthamiana* plants were elicited with 1 µM flg22 (EZBiolabs) or 500 ng/µl (equals ∼1 µM) poly(I:C). As a control, leaf discs were treated with water or treated with PBS. Elicitor and control treatment was performed by addition of the elicitor or the controls to leaf disks acclimated overnight in ultrapure water. After addition of the elicitor, leaf disks were vacuum infiltrated for 10 min. Samples were taken after an additional 20 min of incubation. MPK phosphorylation was determined in protein extracts obtained from elicitor or control-treated leaf disks using immunoblots probed with antibodies against phosphor-p44/42 ERK (Cell Signaling Technology, Beverly, MA, USA) and horseradish peroxidase (HRP)-labeled secondary antibodies for luminescence detection (SuperSignal™ West Femto Maximum Sensitivity Substrate, ThermoFisher).

### Analysis of ROS production

Leaf discs excised from 4-week-old *A. thaliana* or *N. benthamiana* plants were incubated overnight in 96-well plates with 600 μL of deionized, ultra-pure water. The next day deionized, ultra-pure water was replaced with 100 μL reaction solution containing 50 μM luminol and 10 μg/mL horseradish peroxidase (Sigma, USA) together with or without 1μM flg22 or 500 ng/μl polyI:C. Luminescence was determined with a luminometer (BMG LABTECH, FLUOstar®Omega) at 1.5 minute intervals for a period of 40 minutes. Mean values obtained for 10 leaf discs per treatment were expressed as mean relative light units (RLU).

### Seedling growth inhibition assay

Seeds were surface-sterilized and grown vertically at 22°C under 12h/12h light/dark periods in square petri-dishes on half-strength Murashige and Skoog (MS) basal medium (pH 5.8) containing 0.5 g/L MES and 0.8% agar. 7 days old seedlings were transferred into liquid half-strength Murashige and Skoog (MS) medium with or without 500 ng/µl (equals ∼1 µM) poly(I:C) or 1 µM flg22. The effect of treatment on seedling growth was documented on photographs 12 days after treatment and measured with a ruler.

## Funding

Chinese Scholarship Council (PhD fellowship to CH).

Agence National de la Recherche (ANR-13-KBBE-0005-01)

## Author Contributions

Conceptualization: MH

Methodology: CH, JM, EB, ARS, MH

Investigation: CH, ARS

Visualization: CH, ARS, MH

Funding acquisition: MH

Project administration: MH

Supervision: MH

Writing - original draft: MH

Writing – review & editing: CH, ARS, JM, MH

## Data and materials availability

All data are available in the main text or the supplementary materials

## Supplementary Materials

Figs. S1 to S6

Table S1

## Figures

**Fig. S1.**
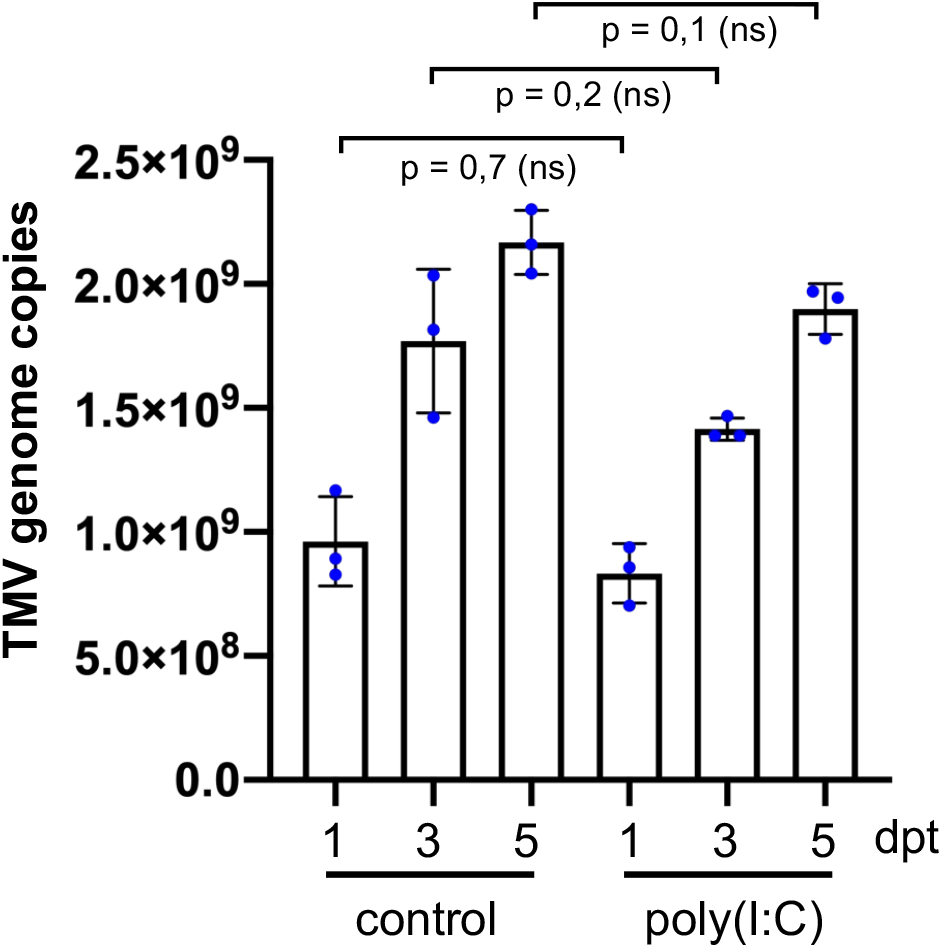
TMV replication is not influenced by poly(I:C). A cell-autonomous (movement-deficient) TMV replicon produces the same number of RNA genome copies in the presence and absence of poly(I:C), as determined by Taqman qPCR. Means of three biological replicates (with SD) per time point and treatment are shown. Two-tailed Mann-Whitney test. dpt, days post transfection.

**Fig. S2.**
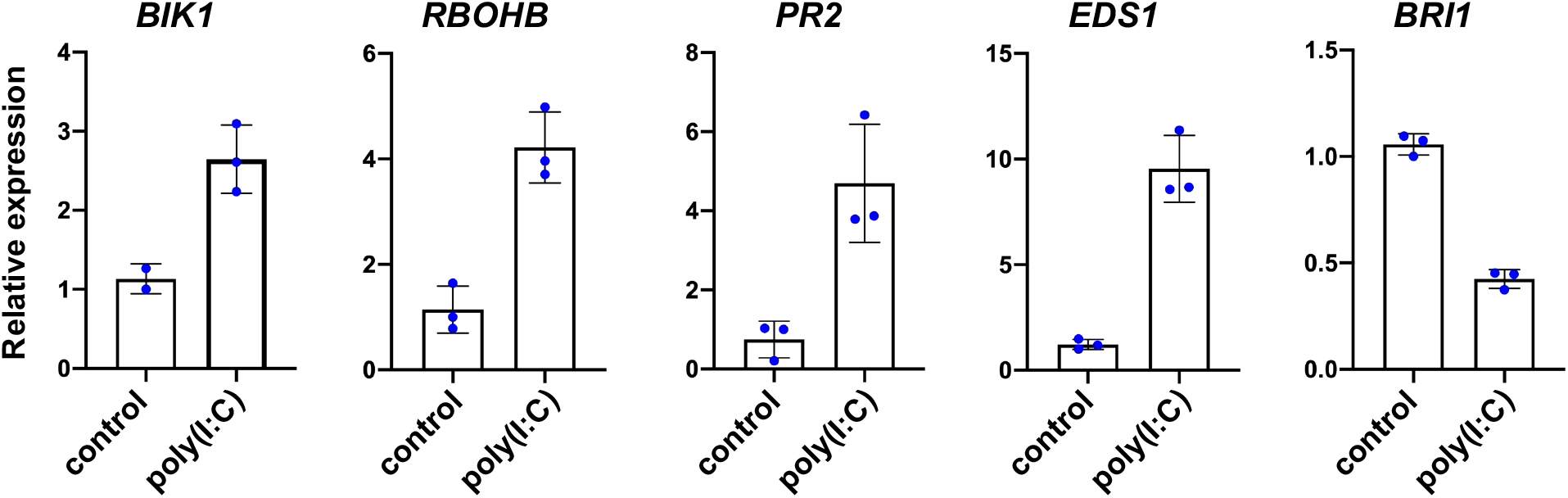
Poly(I:C) induces innate immunity marker genes in *N. benthamiana*. Mean value and SD of gene expression values obtained by RT-qPCR with three biological replicates (blue dots).

**Fig. S3.**
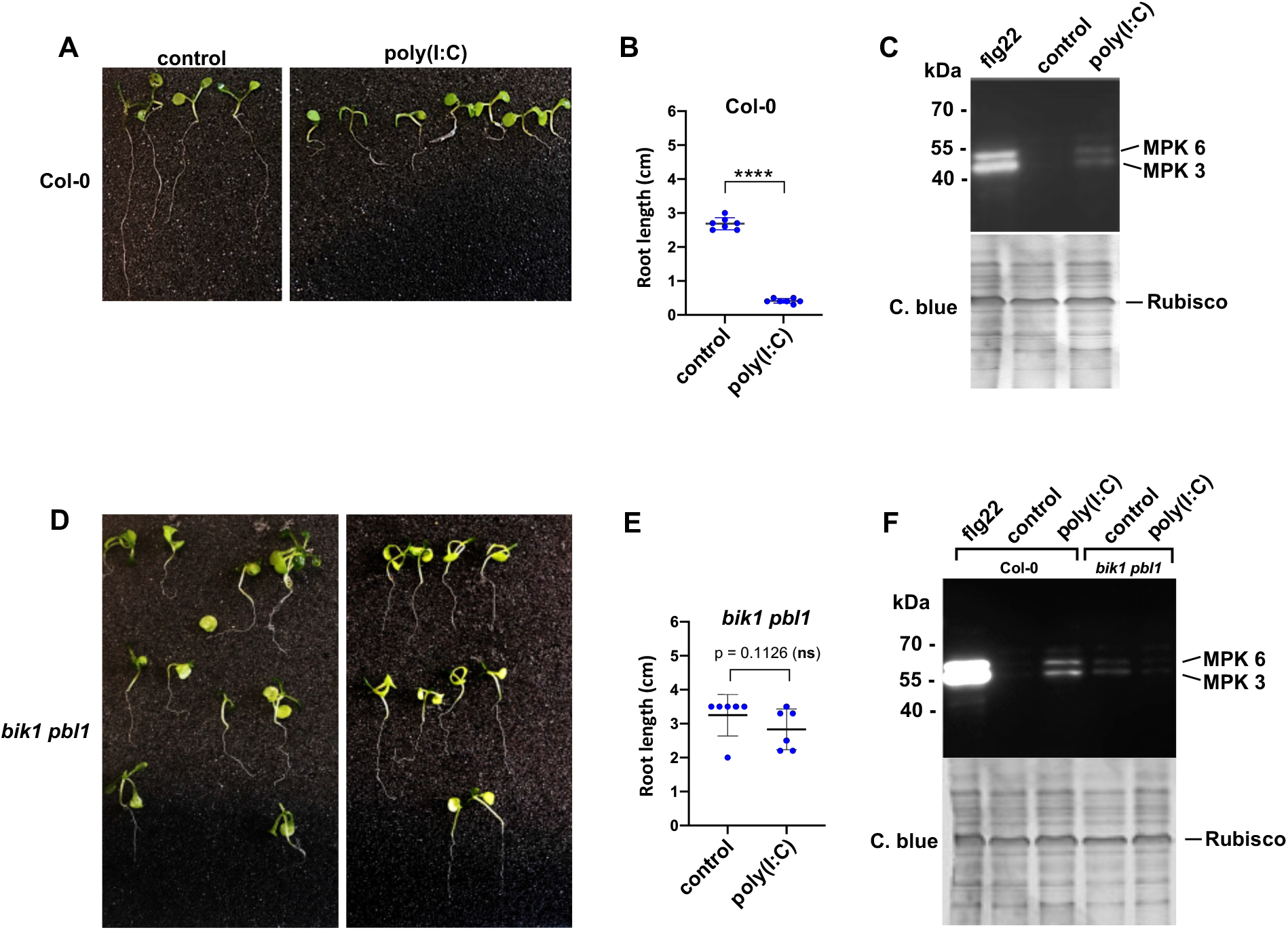
Role of BIK1 in dsRNA-induced root growth inhibition and MPK activation. (**A**) Poly(I:C) inhibits seedling root growth in wild type Arabidopsis wild type Col-0 plants. (**B**) Quantification of dsRNA-induced root growth inhibition in Col-0. Two-tailed Mann-Whitney test; ****, p<0.0001. (**C**) dsRNA-induced MPK activation. Immunoblot detection of phosphorylated MPK. C. blue, Commassie blue-stained gel. (**D**) *bik1 pbl1* plants do not show any significant seedling root growth inhibition in the presence of poly(I:C). (**E**) Quantification of root length inhibition in *bik1 pbl1*. Two-tailed Mann-Whitney test. (**F**) *bik1 pbl1* plants do not show MPK activation in the presence of poly(I:C). Immunoblot detection of phosphorylated MPK. C. blue, Commassie blue-stained gel.

**Fig. S4.**
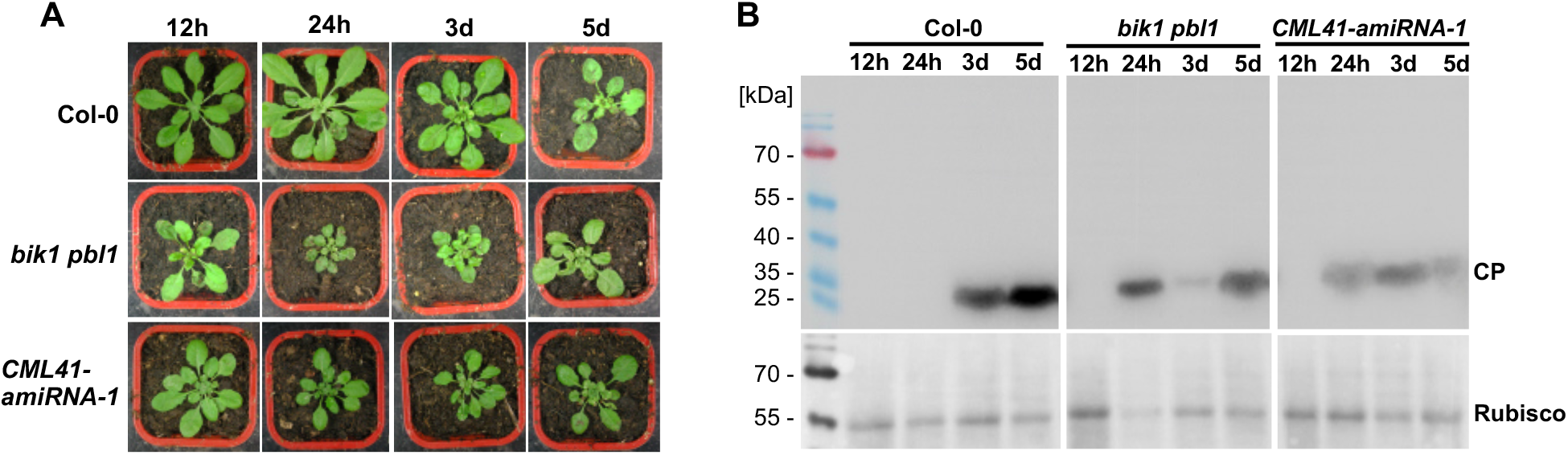
BIK1 and CML41 inhibit systemic virus movement. (**A**) Representative symptom phenotypes at 21 dpi of Arabidopsis Col-0 plants, *bik1 pbl1* mutant plants and plants transgenic for *CML41-amiRNA-1* that were locally inoculated with ORMV and from which the inoculated leaves were removed at the indicated times. Whereas systemic leaves of Col-0 plants show symptoms on plants that carried the inoculated leaves for 3 or more days (d) following inoculation, the systemic leaves of the *bik1 pbl1* mutant and of the CML41-amiRNA expressing plants show symptoms already if the inoculated leaves were present for only 24 hours (h). (**B**) Immunoblot analysis of the youngest systemic leaves at 21 dpi with antibodies against viral coat protein (CP). The pattern of CP expression in the systemic leaves confirms that in wild type Col-0 plants the virus needs between 24 h and 3 d to exit the inoculated leaves and move systemically, whereas the time needed for systemic movement is reduced to less than 24 h in the *bik1 pbl1* mutant and of the CML41-amiRNA expressing plants, thus indicated a role of BIK1 and CML41 in restricting virus movement.

**Fig. S5.**
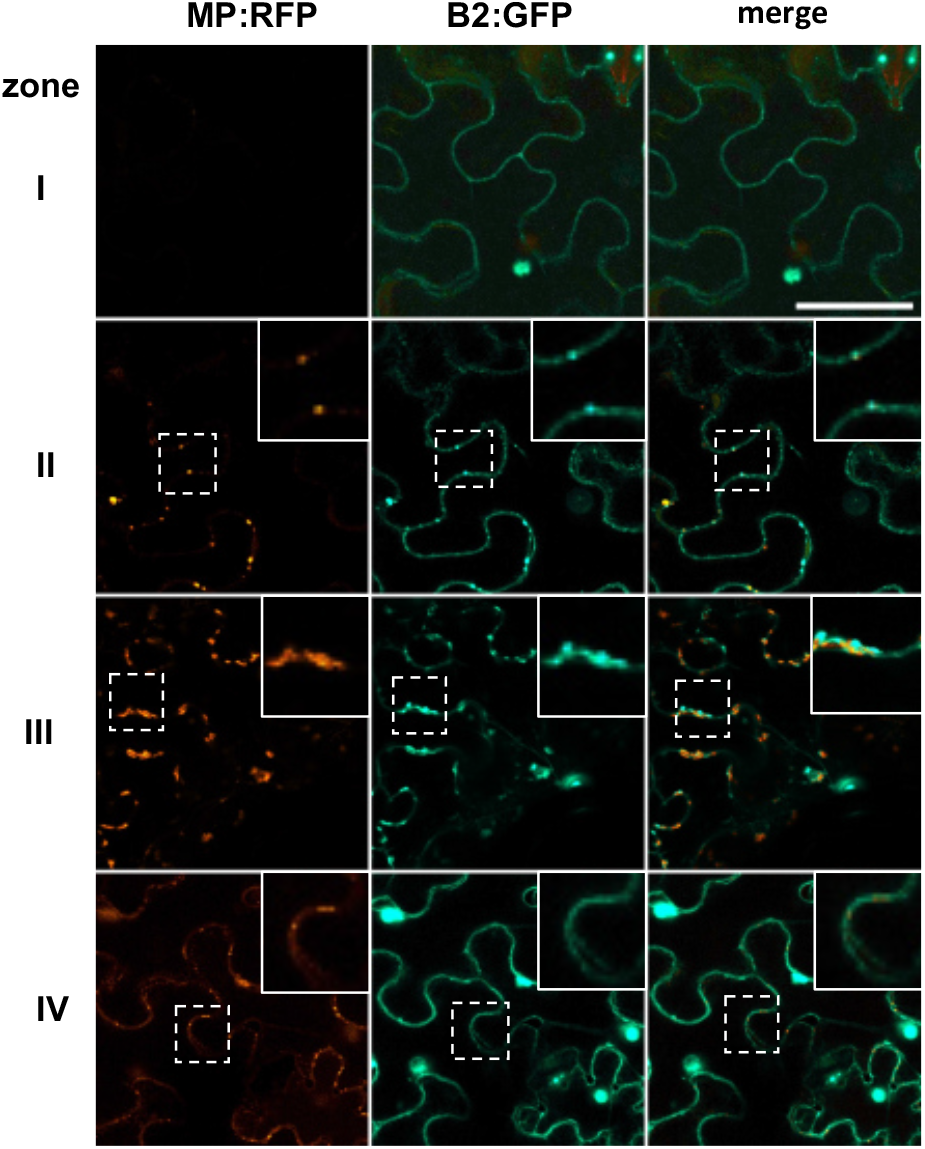
dsRNA accumulation in the different zones of local TMV infection. The MP of TMV-MP:RFP is tagged with RFP and the accumulating dsRNA is imaged through binding of the *Flock house virus* B2 protein fused to GFP (B2:GFP). In cells of zone I (non-infected cells ahead of infection) B2:GFP shows a nucleo-cytoplasmic distribution, which is the typical distribution of this protein in the absence of dsRNA (*31*). In cells at the virus front (zone II), B2:GFP co-localizes to MP:RFP to spots at the cell wall (likely at PD) indicating the localization of early virus-replication complexes (VRCs) engaged in virus replication and virus movement. In zone III, the VRCs have grown in size and accumulate high amounts of dsRNA consistent with high levels of virus replication to produce virus progeny. In zone IV, the MP is no longer expressed. The B2:GP-tagged VRCs now appear rounded and residual MP:RFP is still seen in PD. Inlay images show magnifications of image areas framed by a dashed line. Scale bar = 20 µm.

**Fig. S6.**
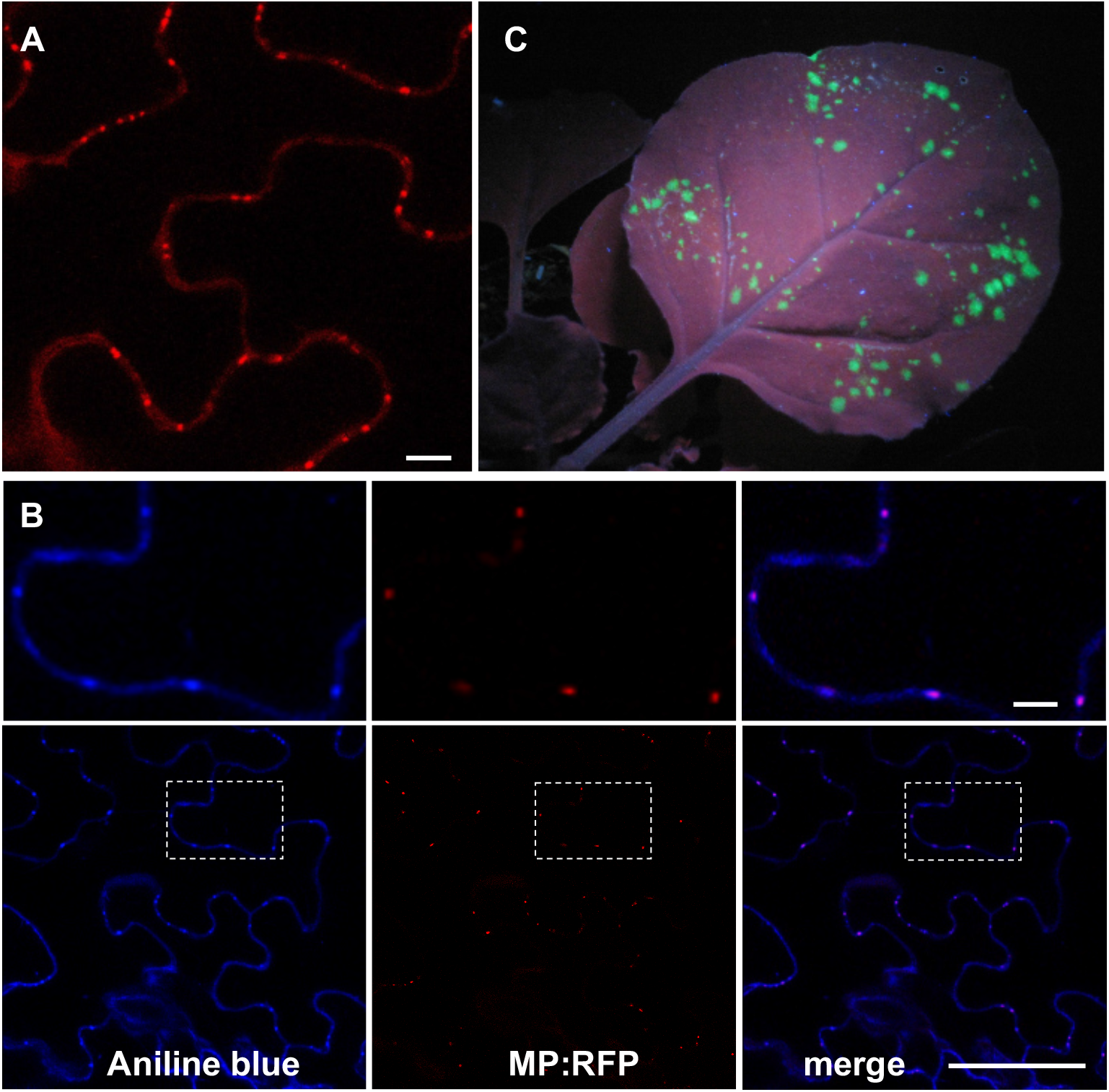
*N. benthamiana* line expressing functional MP:RFP. (**A**) MP:RFP localizes to distinct locations at the plant cell wall. Scale bar, 10 µm. (**B**) The MP:RFP localizes to PD as revealed by callose staining with aniline blue. Scale bar = 100 µm. The upper images show 3.2 x magnifications of the areas framed by a dashed line in the lower images. Scale bar, 10 µm. (**C)** The stably expressed MP:RFP in this line is functional as it complements infection by the MP-deficient TMV construct TMVΔM-GFP (*34*).

**Table S1.**
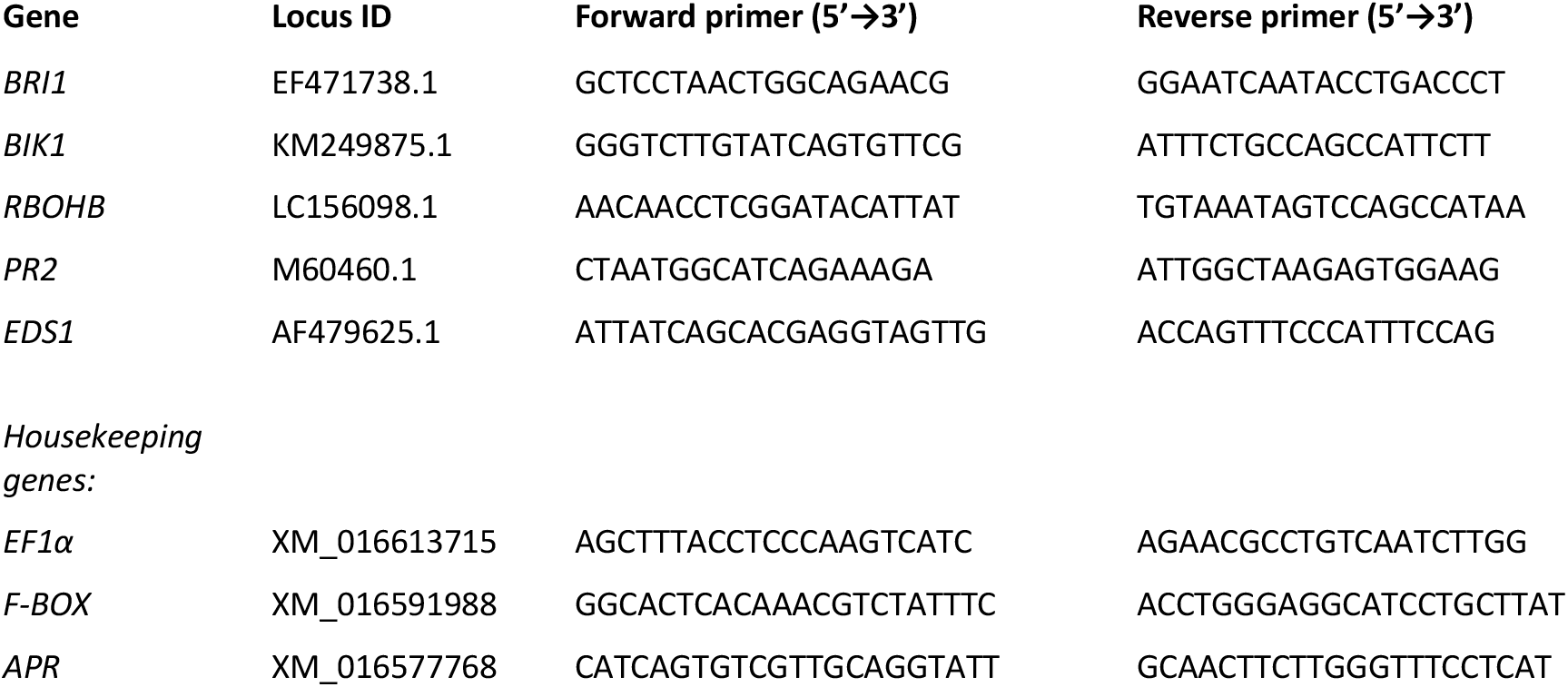
Primers used for RT-qPCR

